# Analysis of Chromatin Data Supports a Role For CD14+ Monocytes/Macrophages in Mediating Genetic Risk for Juvenile Idiopathic Arthritis

**DOI:** 10.1101/2022.07.13.499916

**Authors:** Elizabeth A. Crinzi, Emma K. Haley, Kerry E. Poppenberg, Kaiyu Jiang, Vincent M. Tutino, James N. Jarvis

**Author notes:** Correspondence: James N. Jarvis.

## Abstract

**Introduction:** Genome wide association studies (GWAS) have identified multiple regions that confer genetic risk for the polyarticular/oligoarticular forms of juvenile idiopathic arthritis (JIA). However, genome-wide scans do not identify the cells impacted by genetic polymorphisms on the risk haplotypes or the genes impacted by those variants. We have shown that genetic variants driving JIA risk are likely to affect both innate and adaptive immune functions. We provide additional evidence that JIA risk variants impact innate immunity.

**Materials and Methods:** We queried publicly available H3K4me1/H3K27ac ChIP-seq data in CD14+ monocytes to determine whether the linkage disequilibrium (LD) blocks incorporating the SNPs that tag JIA risk loci showed enrichment for these epigenetic marks. We also queried monocyte/macrophage GROseq data, a functional readout of active enhancers. We defined the topologically associated domains (TADs) encompassing enhancers on the risk haplotypes and identified genes within those TADs expressed in monocytes. We performed ontology analyses of these genes to identify cellular processes that may be impacted by these variants. We also used whole blood RNAseq data from the Genotype-Tissue Expression (GTEx) data base to determine whether SNPs lying within monocyte GROseq peaks influence plausible target genes within the TADs encompassing the JIA risk haplotypes.

**Results:** The LD blocks encompassing the JIA genetic risk regions were enriched for H3K4me1/H3K27ac ChIPseq peaks (p=0.00021 and p=0.022) when compared to genome background. Eleven and sixteen JIA were enriched for resting and activated macrophage GROseq peaks, respectively risk regions (p=0.04385 and p=0.00004). We identified 321 expressed genes within the TADs encompassing the JIA haplotypes in human monocytes. Ontological analysis of these genes showed enrichment for multiple immune functions. Finally, we found that SNPs lying within the GROseq peaks are strongly associated with expression levels of plausible target genes in human whole blood.

**Conclusions:** These findings support the idea that both innate and adaptive immunity are impacted by JIA genetic risk variants.

## Introduction

While multiple genetic risk loci for juvenile idiopathic arthritis (JIA) have been identified and validated,(1-4) the field is still faced with three important tasks: (1) identifying the actual causal variants on the risk haplotypes that exert the biological effects that confer risk; (2) identifying the cells in which those variants exert their effects; and (3) identifying the genes that are impacted by the causal variants (“target genes”).(5) We have recently shown that the latter 2 tasks can be facilitated by understanding the chromatin architecture that encompasses the JIA risk <u>regions.(6-8).</u> For example, the JIA risk haplotypes are highly enriched (compared to genome background) for H3K4me1/H3K27ac ChIPseq peaks, epigenetic signatures typically associated with functional enhancers. This enrichment can be seen in CD4+ T cells, a cell long thought to be involved in JIA pathogenesis (9), and also in neutrophils, for which there is accumulating body of evidence for involvement in JIA pathobiology (10, 11). Furthermore, activation markers of innate immunity, the myeloid related proteins (MRP), remain our most reliable biomarkers of disease activity and durability of remission (12, 13). Thus, we have hypothesized that genetic variants that confer risk for JIA are likely to impact both innate and adaptive immune systems (14).

There is accumulating evidence that other cells of the myeloid lineage, especially CD14+ monocytes/macrophages, play a role in the immunobiology of JIA.(15) We therefore examined the chromatin architecture encompassing the JIA risk <u>regions</u> using publicly available genomic data from CD14+ monocytes and macrophages. Our findings add further support to the notion that risk-driving genetic variants in JIA impact both innate and adaptive immune functions.

## Materials and Methods

Figure 1 provides shows the work flow processes through which we queried the JIA risk regions. Further details are provided in each of the following sections.

**Figure 1.**
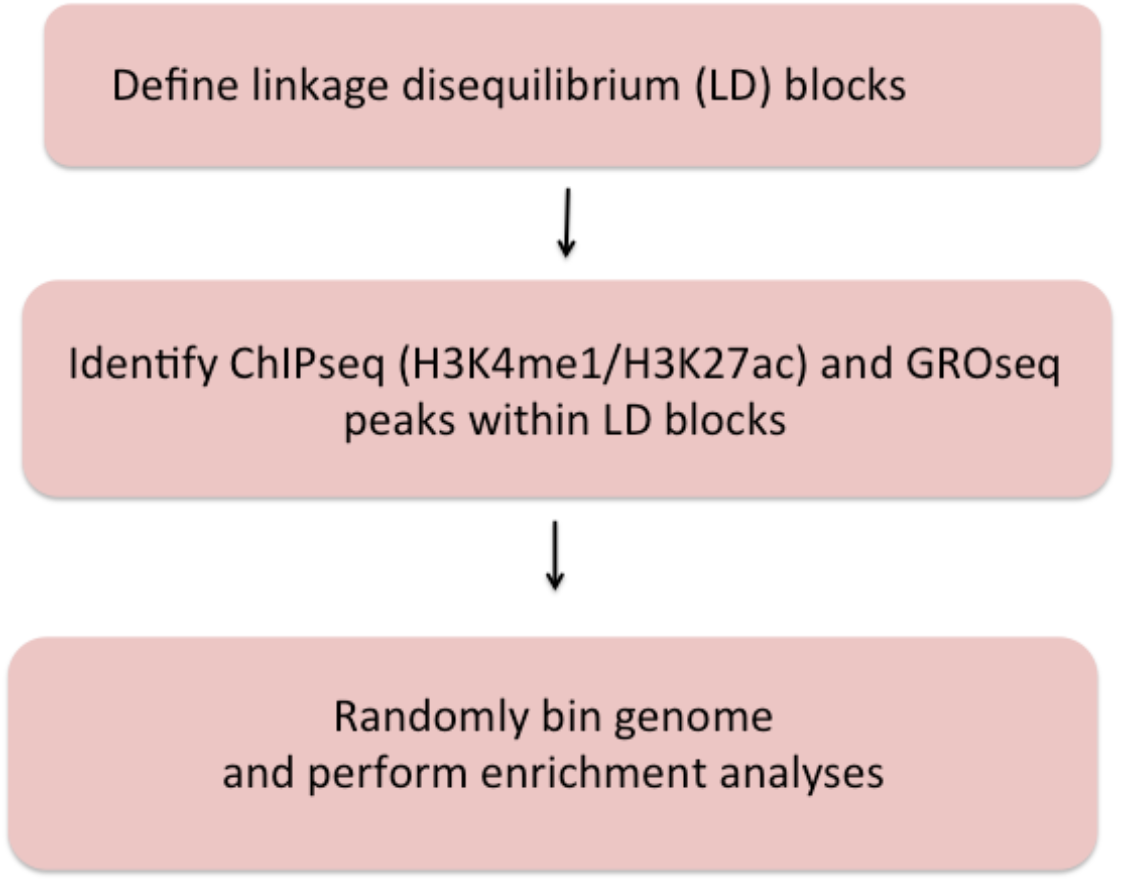
Summary of methods used to query JIA risk regions using the Single Nucleotide Polymorphism Annotator (SNiPA [21]) and publicly available genomic data.

### Disease phenotypes and identification of risk haplotypes

We queried the <u>JIA risk-associated SNPs</u> identified in the Hersh(1) and Herlin(2) reviews, the Hinks Immunochip study (3), and the McIntosh meta-analysis (4) of GWAS data. All of these studies include patients with both oligoarticular JIA and rheumatoid factor-negative, polyarticular JIA. The rationale for this strategy has been that these phenotypes exhibit striking commonalties, including: (1) age of onset typically before puberty; (2) female:male predominance of 3:1; and (3) risk for chronic inflammatory eye disease (uveitis) in younger patients expressing antinuclear antibodies. Although oligoarticular and polyarticular disease are currently categorized as distinct forms of JIA based on the number of involved joints at disease presentation, it is increasingly recognized that this is an arbitrary classification. For example, more than half of children who present with an oligoarticular phenotype pursue a polyarticular disease course (16). In addition, oligo- and polyarticular (RF-negative) subtypes also share HLA associations (17, 18). Thus, the field increasingly views these 2 phenotypes as a disease continuum, sometimes referred to as “polygo” JIA (pJIA) (19, 20), rather than distinct nosocomial entities.

To identify <u>linkage disequilibrium blocks associated with the risk SNPs</u>, we used the Single Nucleotide Polymorphism Annotator (SNiPA) (21), available publicly at https://snipa.helmholtz-muenchen.de/snipa3/. We queried European populations, i.e., the populations represented in the referenced genetic studies, using GRCh37/Ensembl 87 and setting r^2^=0.80. We subsequently converted GRCh37/hg19 coordinates to GRCh38/hg38 using the liftover tool to map relevant chromatin data. These regions are shown in **Table 1**.

**Table 1:** Positional Information of 36 JIA risk haplotypes (r^2^ = 0.80) in hg38

| Locus Name | SNP | LD Block | Length |
| --- | --- | --- | --- |
| PTPN22 | rs6679677 | chr1:113,761,186-113,834,946 | 73,760 |
| ATP882-IL6R | rs11265608 | chr1:154,319,242-154,406,893 | 87,651 |
| STAT4 | rs10174238 | chr2:191,079,016-191,108,308 | 29,292 |
| IL2-21 | rs1479924 | chr4:122,151,854-122,619,603 | 467,749 |
| ANKRD55 | rs71624119 | chr5:56,141,024-56,146,422 | 5,398 |
| ERAP2-LNPEP | rs27290 | chr5:96,884,383-97,038,046 | 153,663 |
| C5orf56-IRF1 | rs4705862 | chr5:132,477,527-132,496,822 | 19,295 |
| HLA-DQB1-DQA2 | rs7775055 | chr6:32,422,420-32,712,215 | 289,795 |
| IL2RA | rs7909519 | chr10:6,028,313-6,055,320 | 27,007 |
| SH2B3-ATXN2 | rs3184504 | chr12:111,395,984-111,645,358 | 249,374 |
| ZFP36L1 | rs12434551 | chr14:68,784,174-68,794,755 | 10,581 |
| PTPN2 | rs2847293 | chr18:12,774,327-12,809,341 | 35,014 |
| TYK2 | rs34536443 | chr19:10,317,045-10,381,598 | 64,553 |
| RUNX1 | rs9979383 | chr21:35,323,611-35,365,944 | 42,333 |
| UBE2L3 | rs2266959 | chr22:21,556,931-21,628,971 | 72,040 |
| IL2RB | rs2284033 | chr22:37,135,077-37,141,474 | 6,397 |
| TIMMDC1-CD80 | rs4688011 | chr3:119,406,355-119,529,051 | 122,696 |
| JAK1 | rs10889504 | chr1:64,924,820-64,975,081 | 50,261 |
| PRR9-LOR | rs873234 | chr1:153,249,128-153,270,022 | 20,894 |
| PTH1R | rs1138518 | chr3:46,889,988-46,932,682 | 42,694 |
| ILDR1-CD86 | rs111700762 | chr3:122,023,892-122,102,059 | 78,167 |
| LINC-00951 | rs10807228 | chr6:40,198,646-40,261,171 | 62,525 |
| AHI1-LINC-00271 | rs9321502 | chr6:135,303,673-135,371,709 | 68,036 |
| HBP1 | rs111865019 | chr7:107,156,092-107,390,877 | 234,785 |
| WDFY4 | rs1904603 | chr10:48,776,493-48,805,795 | 29,302 |
| RNF215 | rs5753109 | chr22:30,289,571-30,403,731 | 114,160 |
| LTBR | rs2364480 | chr12:6,384,185-6,402,830 | 18,645 |
| IL6 | rs7808122 | chr7:22,729,088-22,771,765 | 42,677 |
| COG6 | rs7993214 | chr13:39,726,191-39,794,464 | 68,273 |
| 13q14 | rs34132030 | chr13:42,481,900-42,492,387 | 10,487 |
| CCR1-CCR3 | rs79893749 | chr3:46,141,688-46,420,292 | 278,604 |
| PRRL5 | rs4755450 | chr11:36,314,713-36,354,471 | 39,758 |
| PRM1-RM12 | rs66718203 | chr16:11,278,799-11,352,790 | 73,991 |
| RUNX3 | rs4648881 | chr1:24,870,664-24,884,324 | 13,660 |
| JAZF1 | rs10280937 | chr7:28,114,765-28,207,370 | 92,605 |
| AFF3-LONRF2 | rs6740838 | chr2:100,196,869-100,221,105 | 24,236 |

### Enrichment analysis: H3K4me1/H3K27ac ChIPseq data

We investigated the presence of potential enhancers within the <u>JIA risk regions by</u> defining enrichment (compared to genome background) for H3K4me1 and H3K27ac histone marks from publicly available ChIPseq data. We used the BedTools software to assess for enrichment in CD14+ human monocytes, following the method described by Poppenberg et al.(22) Briefly, we mined publicly available CD14+ H3K4me1 and H3K27ac ChIPseq using the Cistrome database (http://cistrome.org/db/#/ ). We first queried a single data set for each H3K4me1 (GEO accession #GSM1003535) and H3K27ac (GEO accession #GSM2818004). We then independently validated these findings from 3 additional data sets for H3K4me1 (GSM1102793, GSM995353, and GSM1003535) and H3K27ac (GSM1102782, GSM995355, and GSM995356). Using the intersect command in BedTools, we overlapped these peaks with the JIA haplotypes. We created 36 random regions of 86,676 base pairs, or the average length of all the JIA haplotypes, to compare ChIPseq peak enrichment against the background genome. We repeated this random region generation 1000 times to approximate a normal distribution. We then calculated the associated *p* value with this normal distribution by aligning the curve with number of overlaps between the JIA haplotypes and the ChIP-seq peaks. We considered a *p* value of *p*<0.05 as statistically significant.

### Enrichment analysis: GROseq data

Although H3K4me1/H3K27ac ChIPseq peaks are epigenetic characteristics associated with enhancers, additional functional data can also be used to identify these regulatory elements. Danko et al(23) have shown that global run-on sequencing (GROseq) data can be used to identify the bi-directional RNA synthesis that is a hallmark of functional enhancers. We therefore use dReg(23) to identify GROseq peaks using publicly available data from resting macrophages and macrophages activated <u>with Kdo2-Lipid-A (KLA), a lipopolysaccharide derived from E. coli.(24)</u> Using the intersect command in BedTools, we overlapped these GROseq peaks with the JIA haplotypes. We created 36 random regions of the average length of all the haplotypes, to compare the GROseq enrichment against the background genome. We repeated this random region generation 1000 times to approximate a normal distribution. We then calculated the associated *p* value with this normal distribution by aligning the curve with number of overlaps between the JIA haplotypes and the GROseq peaks. We considered a *p* value of *p*<0.05 as statistically significant.

### Defining topologically associated domains (TADs)

Identifying target genes influenced by enhancers that harbor disease-driving genetic variants can be facilitated by knowing the larger 3D chromatin architecture that encompasses the risk haplotypes that harbor the relevant enhancers and variants (8). For example, Gasperini et al (25) showed that, on a genome-wide CRISPRi screen, 71% of enhancers regulate genes within the same TAD. We therefore adapted the method described by Poppenberg et al (22) to identify TAD structures that encompass the JIA risk haplotypes in monocytes. In brief, we first analyzed a publicly available HiC data set (26) using Juicebox software (27) following the method described by Kessler et al (8) to identify interacting regions in the monocyte THP-1 cell line. Since TADs are anchored by complexes of CTCF and cohesion (28), we sought to assure the relevance of these analyses, excluding from subsequent analysis any HiC-defined interacting region that did not also have clear CTCF anchors as identified from a publicly available CTCF ChIPseq data set generated from the primary human monocytes (29) (**Figure 2**).

**Fig. 2.**
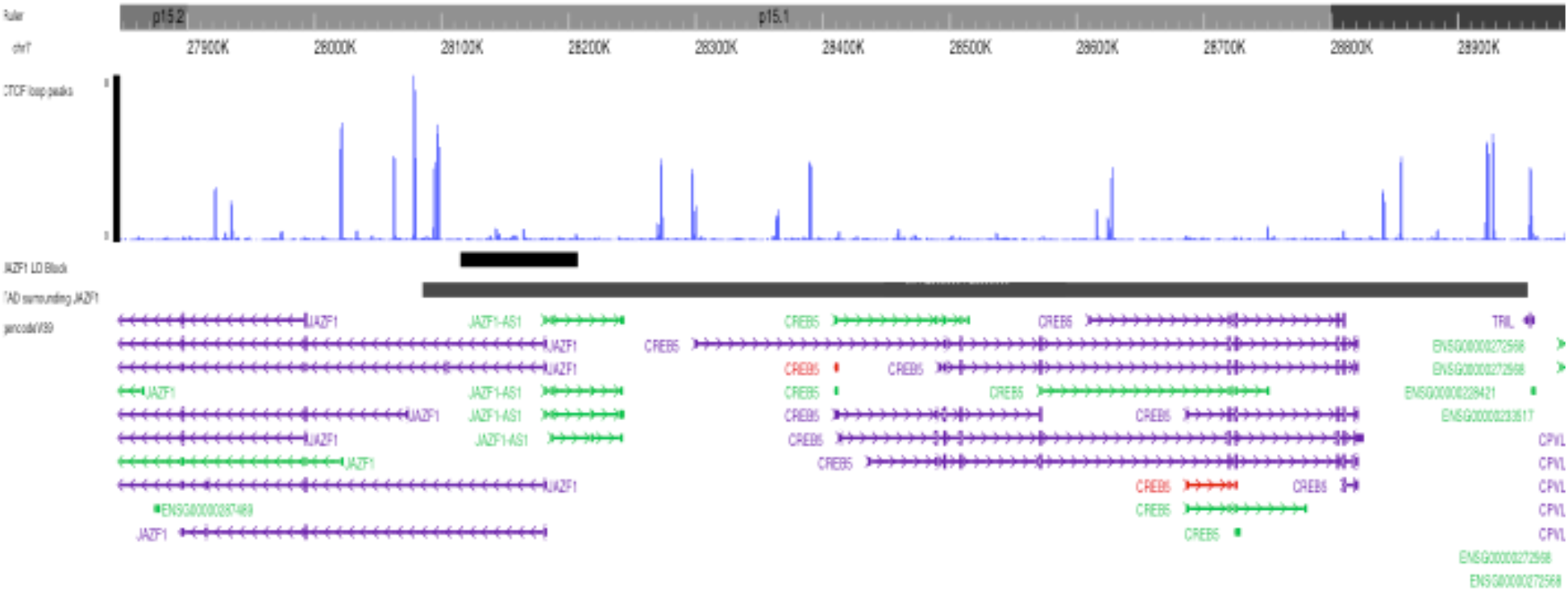
Washington University Epigenome Browser visualization of the *JAZF1* risk locus (short solid bar), the corresponding TAD that surrounds the *JAZF1* locus (longer solid bar) and, CTCF ChIPseq peaks (blue) and the genes located within the TAD. Note that each end of the TAD is anchored by CTCF.

### Identification and ontology analysis of expressed genes within the HiC/CTCF ChIPseq-defined TADs

To characterize the most likely gene targets of identified enhancers, we used the UCSC and Washington University Genome Browsers to align each defined TAD with RefSeq genes as shown in **Figure 2**. Pseudogenes and non-coding RNA were filtered out within the browser settings. RefSeq genes were filtered by expression level in human monocytes using RNAseq data published by Schulert et al (30). We excluded any gene with a TPM of ≤ 1. We determined functionality of the expressed genes using gene ontology analysis. We used the publicly available Gene Ontology enRIchment anaLysis and visuaLizAtion (GORILLA) to compare the significance of the expression of these genes as compared to a background set comprising all genes expressed in human monocytes. We input the significant GO terms from GORILLA into the Reduce Visualize Gene Ontology (REVIGO) tool to visualize gene functionalities by semantic categories.

### Enrichment analysis: identifying enhancers that harbor expression-altering variants identified using a massively parallel reporter assay

In myeloid K562 cells, we used a massively parallel reporter assay (MPRA) identical to that described by Tewhey et al (31) to identify SNPs that have the intrinsic ability to alter gene expression when compared to the common allele (32). The assay uses bar coded oligonucleotides (“oligos”) with a minimal promoter driving green fluorescence protein expression, making it suitable for querying allelic effects on non-coding genomic functions. The allele of interest is placed in the center of the 180 bp oligo with the cognate flanking sequences on either side, providing a degree of genomic context for the reporter. Bar coding allows expression levels (determined by RNAseq) to be determined for each SNP and compared to the common allele. For these assays, we studied both resting K562 cells and cells stimulated with interferon gamma (K562+IFNG) (32) a pathologically relevant ligand (33).

Using the ‘intersect’ command in BedTools, we overlapped these GROseq peaks with the MPRA regions. We created 18 random regions of the average length of all the MPRA regions, to compare the GROseq enrichment against the background genome. We repeated this random region generation 1000 times to approximate a normal distribution. We then calculated the associated *p* value with this normal distribution by aligning the curve with number of overlaps between the MPRA regions and the GROseq peaks. We considered a *p* value of *p*<0.05 as statistically significant.

### Linking genetic variants in GROseq peaks to human expression whole blood expression levels

As proof of concept that SNPs within enhancers might influence gene expression, we used the data from the analysis of MPRA SNPs and GROseq peaks as described above to determine whether the genetic variants within the GROseq peaks could be demonstrated to influence gene expression in human peripheral blood cells. As noted, the MPRA was performed in myeloid K562 cells, which have similar TAD features to myeloid THP-1 cells and, thus, to human CD14+ monocytes.(8) From the TAD data, we then identified genes that were likely to be regulated by the GROseq-defined enhancers, and thus be influenced by the MPRA-identified SNPs. We used the expression quantitative trait locus (eQTL) calculator on the Gene-Tissue Expression (GTEx) web site [https://www.gtexportal.org/home/testyourown] to query the effects of individual SNPs on gene expression derived from 670 individuals in the GTEx project. The eQTL software calculates statistical significance after correcting for multiple comparisons and the threshold for significance varies with individual genes and SNPs.

## Results

### Enrichment analysis: H3K4me1/H3K27ac ChIPseq data

We investigated the enrichment of H3K4me1 and H3K27ac ChIPseq peaks within 36 <u>LD blocks conferring</u> JIA risk in CD14+ monocytes to determine whether these regions are enriched for epigenetic signatures of enhancers, as they are in neutrophils and CD4+ T cells.(6, 7) Out of the 36 risk haplotypes, 22 were enriched for both H3K4me1 and H3K27ac peaks, while 7 haplotype regions were enriched for H3K4me1 alone. Thus, 7 of 36 haplotypes did not contain any overlaps with ChIP-seq peaks for either H3K4me1 or H3K27ac. This was expected, as it is unlikely that genetic risk in every region operates in every relevant cell type. The presence or absence of histone ChIPseq peaks for each haplotype is outlined in **Table 2**. Overall, the JIA haplotypes were enriched in CD14+ monocytes at to a significant degree: *p*<0.022 for H3K27ac enrichment and *p*<0.00021 for H3K4me1. We were able to independently confirm these results by performing the enrichment analyses on 3 independent data sets for H3K4me1 with significant p-values for each data set. (GSM1102793 p=0.0012; GSM995353 p=0.0012; and GSM1003535 p<0.00021). We were also able to independently validate the findings from H3K27ac ChIPseq enrichment analysis (GSM1102782 p=0.00014; GSM995355 p=0.00011); and GSM995356 p=0.00030).

**Table 2.**
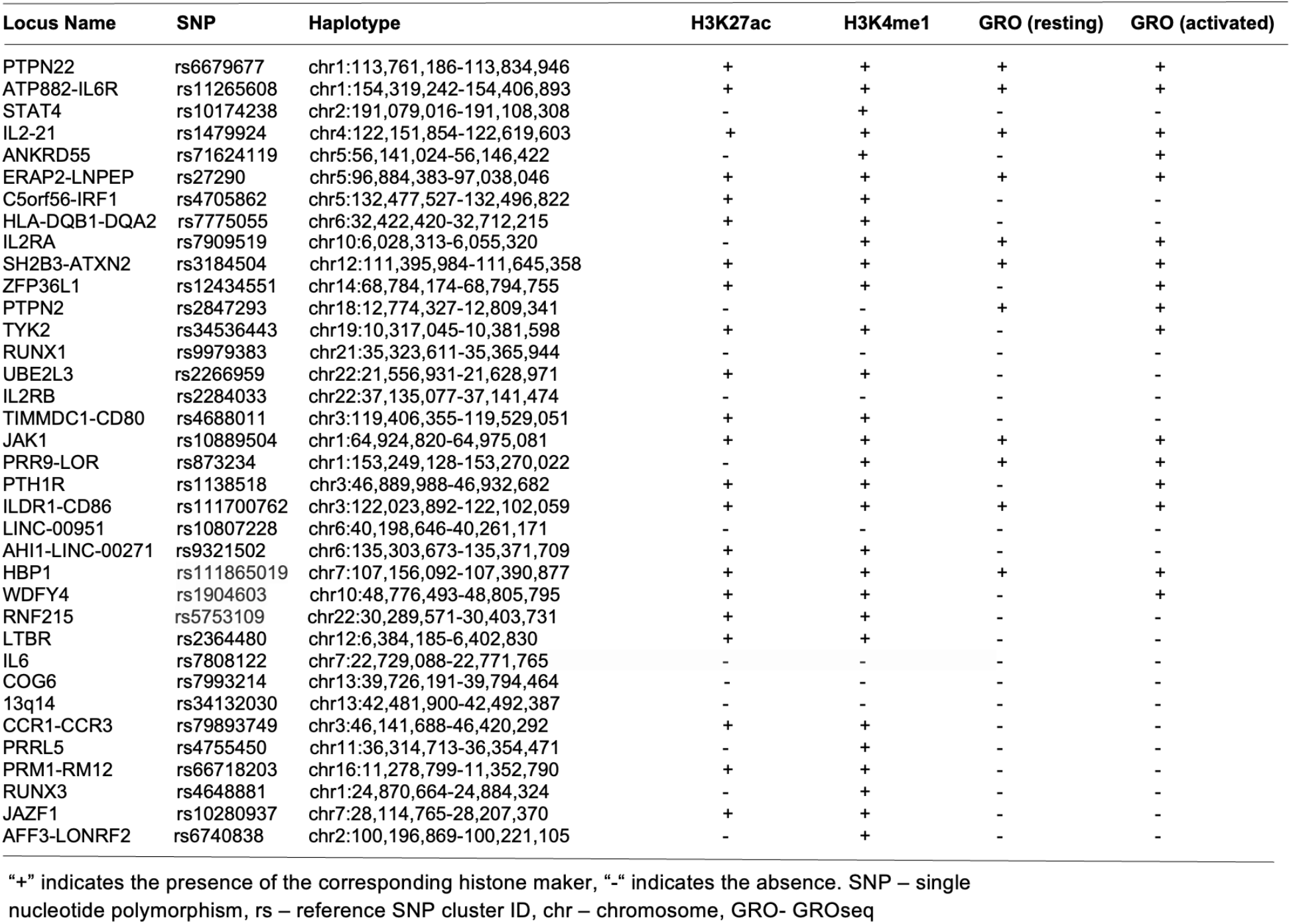
Histone marks present in JIA associated haplotypes

### Enrichment analysis: GROseq data

We also interrogated the JIA risk haplotypes for GROseq peaks in both resting macrophages and macrophages activated by KLA from data published by Kaikkonen et al.(24) We also sought to determine whether the presence of GROseq peaks occurred at a greater-than-expected frequency on JIA haplotypes compared with randomly-selected regions of the genome. There were GROseq peaks in 11 out of the 36 <u>LD blocks</u> in resting macrophages, and in 16 haplotypes in KLA-stimulated macrophages. This pattern occurred at a higher-than-expected frequency using genome background as a comparison in both macrophages (p=0.04385) and in KLA stimulated macrophages (p=0.00004). The five additional <u>LD blocks</u> that became enriched for GROseq peaks after macrophage differentiation with KLA were *ANKRD55, TYK2, ZFP36L1, PTH1R*, and *WDFY4*. In resting macrophages, 7 <u>LD blocks</u> were enriched for both GROseq peaks and H3K27ac peaks, and in 13 haplotypes in KLA-stimulated macrophages (**Table 2**).

### Monocyte GROseq peaks intersect with SNPs identified on MPRA

We next asked whether the GROseq peaks overlapped with expression-altering SNPs that we have identified using an MPRA in myeloid K562cells (32) as described in the *Methods* section. From the MPRA data, we identified 18 chromatin regions containing 42 SNPs lying within H3K4me1/H3K27ac-marked regions derived from ChIPseq data in human CD14+ monocytes. **Table 3** provides chromatin coordinates for these regions. We next sought to determine how many of these regions also encompassed a GROseq peak.

**Table 3:**
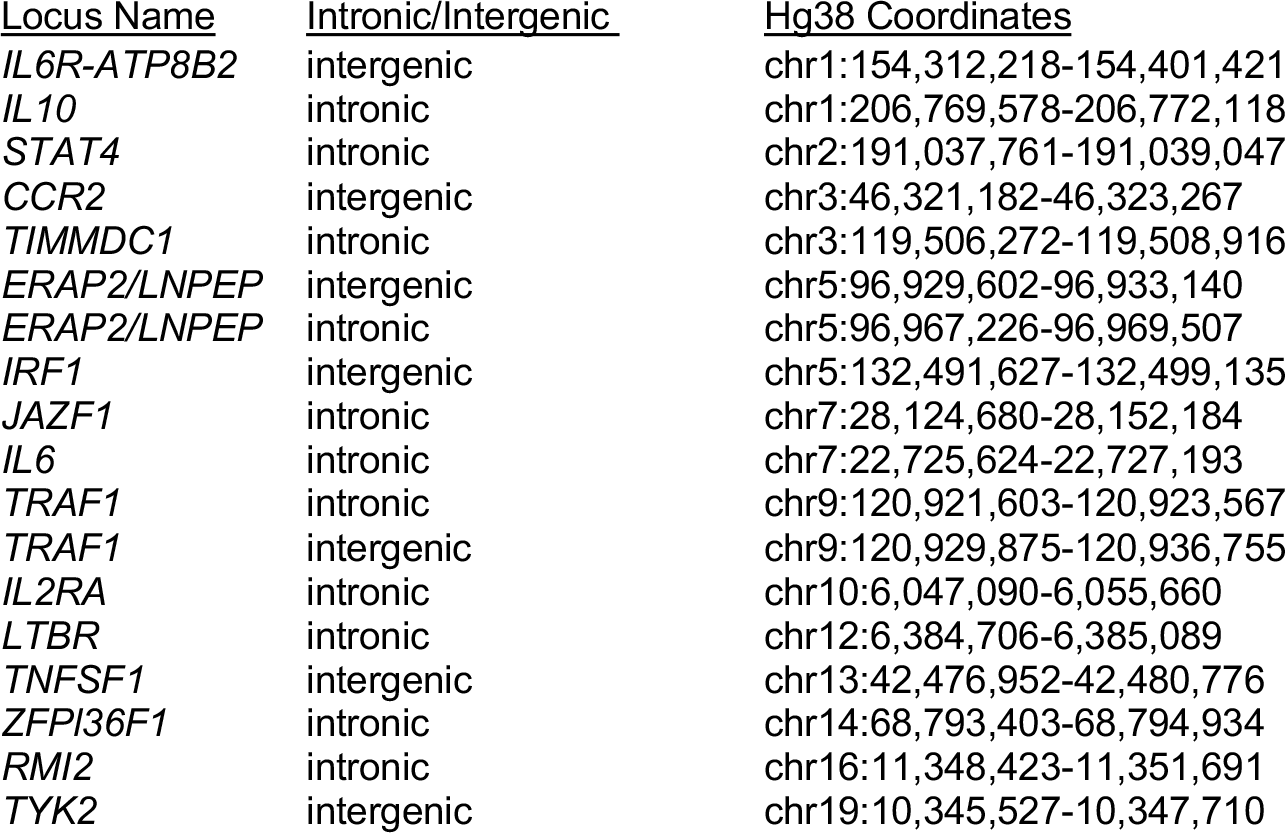
Chromatin coordinates of enhancers identified by MPRA in unstimulated K562 in Hg38

Note that many of these regions are longer that the typical 800-1,000 bp length of individual enhancers. These are broad regions of open chromatin where multiple H3K4me1/H3K27ac peaks are identified, and likely represented so-called super enhancer complexes. We identified an 19 additional SNPs in IFNG-stimulated K562 cells, also lying within H3K4me1/H3K27ac-marked regions (again using human CD14+ monocyte ChIPseq data). **Table 4** provides chromatin coordinates for these regions.

**Table 4:**
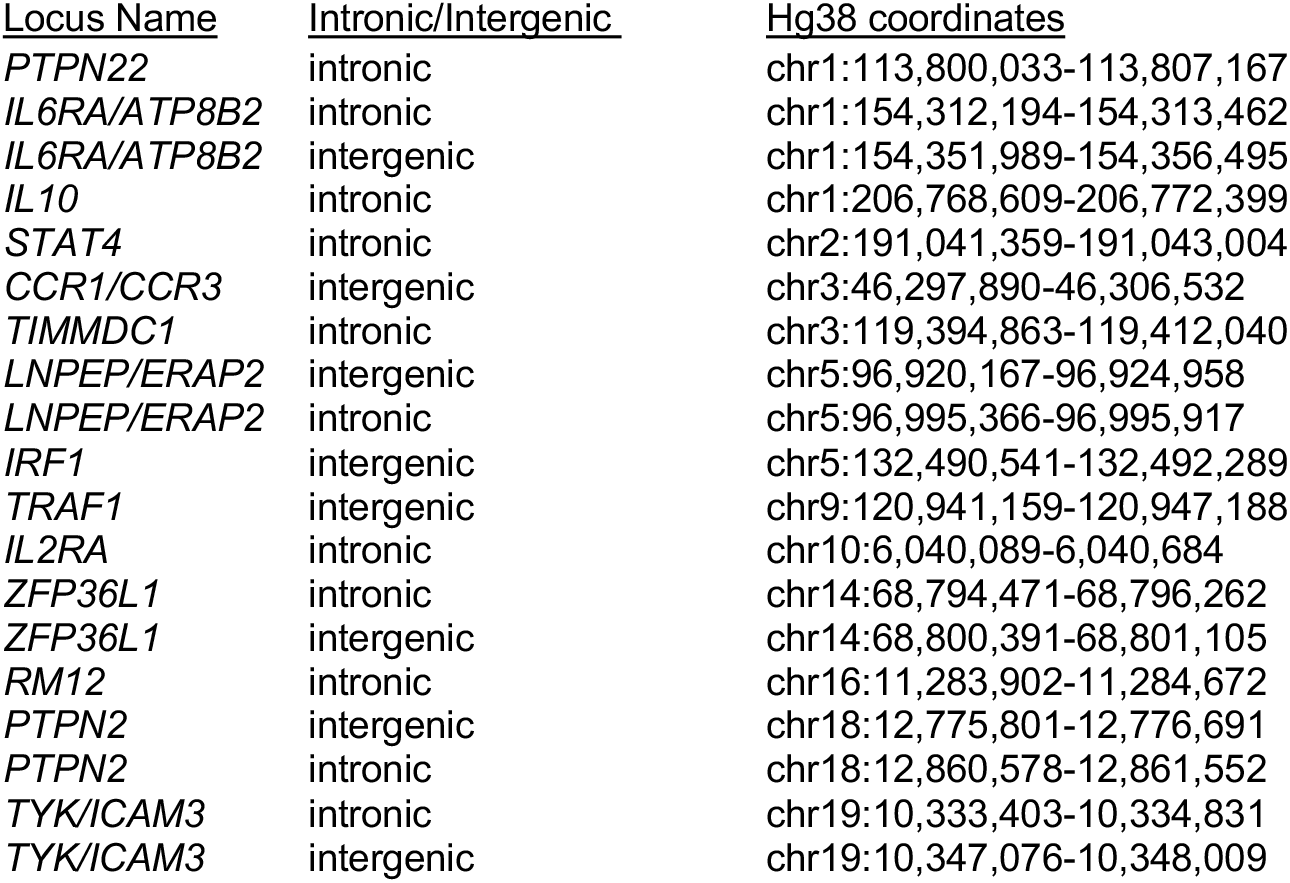
Chromatin coordinates of enhancers identified by MPRA in K562+IFNG in Hg38

We next sought to determine how many of these SNPs also intersect with a GROseq peak. Of the 18 MPRA-identified regions in resting K562 cells, 7 SNPs overlapped GROseq peaks in resting macrophages, and 8 overlapped with GROseq peaks in KLA-stimulated macrophages (**Table 5)**.

**Table 5:** GROseq peaks that overlap SNPs detected on MPRA in resting K562 cells

| <u>Locus Name</u> | <u>Intronic/Intergenic</u> | <u>Hg38 Haplotype</u> | <u>GRO (resting)</u> | <u>GRO (activated)</u> |
| --- | --- | --- | --- | --- |
| *IL6R-ATP8B2* | intergenic | chr1:154,312,218-154,401,421 | + | + |
| *IL10* | intronic | chr1:206,769,578-206,772,118 | - | - |
| *STAT4* | intronic | chr2:191,037,761-191,039,047 | - | - |
| *CCR2* | intergenic | chr3:46,321,182-46,323,267 | + | + |
| *TIMMDC1* | intronic | chr3:119,506,272-119,508,916 | - | - |
| *ERAP2/LNPEP* | intergenic | chr5:96,929,602-96,933,140 | - | - |
| *ERAP2/LNPEP* | intronic | chr5:96,967,226-96,969,507 | - | - |
| *IRF1* | intergenic | chr5:132,491,627-132,499,135 | + | + |
| *JAZF1* | intronic | chr7:28,124,680-28,152,184 | + | + |
| *IL6* | intronic | chr7:22,725,624-22,727,193 | - | - |
| *TRAF1* | intronic | chr9:120,921,603-120,923,567 | + | + |
| *TRAF1* | intergenic | chr9:120,929,875-120,936,755 | - | + |
| *IL2RA* | intronic | chr10:6,047,090-6,055,660 | - | - |
| *LTBR* | intronic | chr12:6,384,706-6,385,089 | - | - |
| *TNFSF1* | intergenic | chr13:42,476,952-42,480,776 | - | - |
| *ZFPI36F1* | intronic | chr14:68,793,403-68,794,934 | - | - |
| *RMI2* | intronic | chr16:11,348,423-11,351,691 | + | + |
| *TYK2* | intergenic | chr19:10,345,527-10,347,710 | + | + |

"+" indicates the presence of a GROseq peak, "-" indicates the absence. chr-chromosome, GRO- GROseq

<u>Out of the 19 MPRA-identified SNPs in K562+IFNG cells, 8 SNPs overlapped GROseq peaks in resting macrophages, and 7 overlapped with GROseq peaks in KLA-stimulated macrophages</u> (Table 6).

**Table 6:** GROseq peaks that overlap SNPs detected on MPRA in K562+IFNG

| <u>Locus Name</u> | <u>Intronic/Intergenic</u> | <u>Hg38 Haplotype</u> | <u>GRO (resting)</u> | <u>GRO (activated)</u> |
| --- | --- | --- | --- | --- |
| *PTPN22* | intronic | chr1:113,800,033-113,807,167 | - | - |
| *IL6RA/ATP8B2* | intronic | chr1:154,312,194-154,313,462 | + | + |
| *IL6RA/ATP8B2* | intergenic | chr1:154,351,989-154,356,495 | + | + |
| *IL10* | intronic | chr1:206,768,609-206,772,399 | - | - |
| *STAT4* | intronic | chr2:191,041,359-191,043,004 | - | - |
| *CCR1/CCR3* | intergenic | chr3:46,297,890-46,306,532 | - | - |
| *TIMMDC1* | intronic | chr3:119,394,863-119,412,040 | + | + |
| *LNPEP/ERAP2* | intergenic | chr5:96,920,167-96,924,958 | - | - |
| *LNPEP/ERAP2* | intronic | chr5:96,995,366-96,995,917 | - | - |
| *IRF1* | intergenic | chr5:132,490,541-132,492,289 | + | - |
| *TRAF1* | intergenic | chr9:120,941,159-120,947,188 | + | + |
| *IL2RA* | intronic | chr10:6,040,089-6,040,684 | - | - |
| *ZFP36L1* | intronic | chr14:68,794,471-68,796,262 | - | - |
| *ZFP36L1* | intergenic | chr14:68,800,391-68,801,105 | - | - |
| *RM12* | intronic | chr16:11,283,902-11,284,672 | - | - |
| *PTPN2* | intergenic | chr18:12,775,801-12,776,691 | - | - |
| *PTPN2* | intronic | chr18:12,860,578-12,861,552 | + | + |
| *TYK/ICAM3* | intronic | chr19:10,333,403-10,334,831 | + | + |
| *TYK/ICAM3* | intergenic | chr19:10,347,076-10,348,009 | + | + |

“+” indicates the presence of a GROseq peak, “-” indicates the absence. chr-chromosome, GRO- GROseq

### Defining topologically associated domains (TADs) and identifying expressed genes within the TADs

As described previously, target genes of enhancers are almost always located within the associated topologically associated domains (TADs) that encompass those enhancers.(25) Therefore, we defined the TADs encompassing the identified enhancers in the <u>JIA LD blocks</u> to begin our investigation of each enhancer’s target genes. We outlined TADs using publicly available Hi-C data and Juicebox software(8, 27) for each of the 29 <u>LD blocks</u> that displayed enrichment for either both H3K27ac and H3K4me1 or H3K4me1 alone. TADs are invariably flanked by CTCF anchors at each end of the loop, so to strengthen the pathological relevance of these analyses, we excluded regions that displayed interactions in HiC data in THP-1 cells that did not also have clear CTCF anchors at either end of the loop in human CTCF ChIPseq data, as described in the *Methods* section and shown in **Figure 2**. We excluded a single region, *PTPN22*, based on these criteria. TADs encompassing all 28 of the enhancer regions are shown in the supplemental results.

Within each of the 28 analyzed TADs, we identified the monocyte genes most likely to be regulated by the enhancers on the JIA risk haplotypes using the UCSC browser and Refseq gene alignment feature. Filtering this list for genes expressed in monocytes with a TPM of >1 using control monocyte expression data from Schulert et al(30) revealed 321 potential target genes. We completed an ontological analysis to compare the enriched (p<0.01) functions of target genes regulated by enhancers on the risk haplotypes. We identified 14 associated biological processes involved in multiple immune related functions (cytokine binding, interleukin-6 receptor binding, and MHC class II receptor activity, etc), as well as other basic cellular processes such as peptide receptor activity (**Figure 3**).

**Fig. 3.**
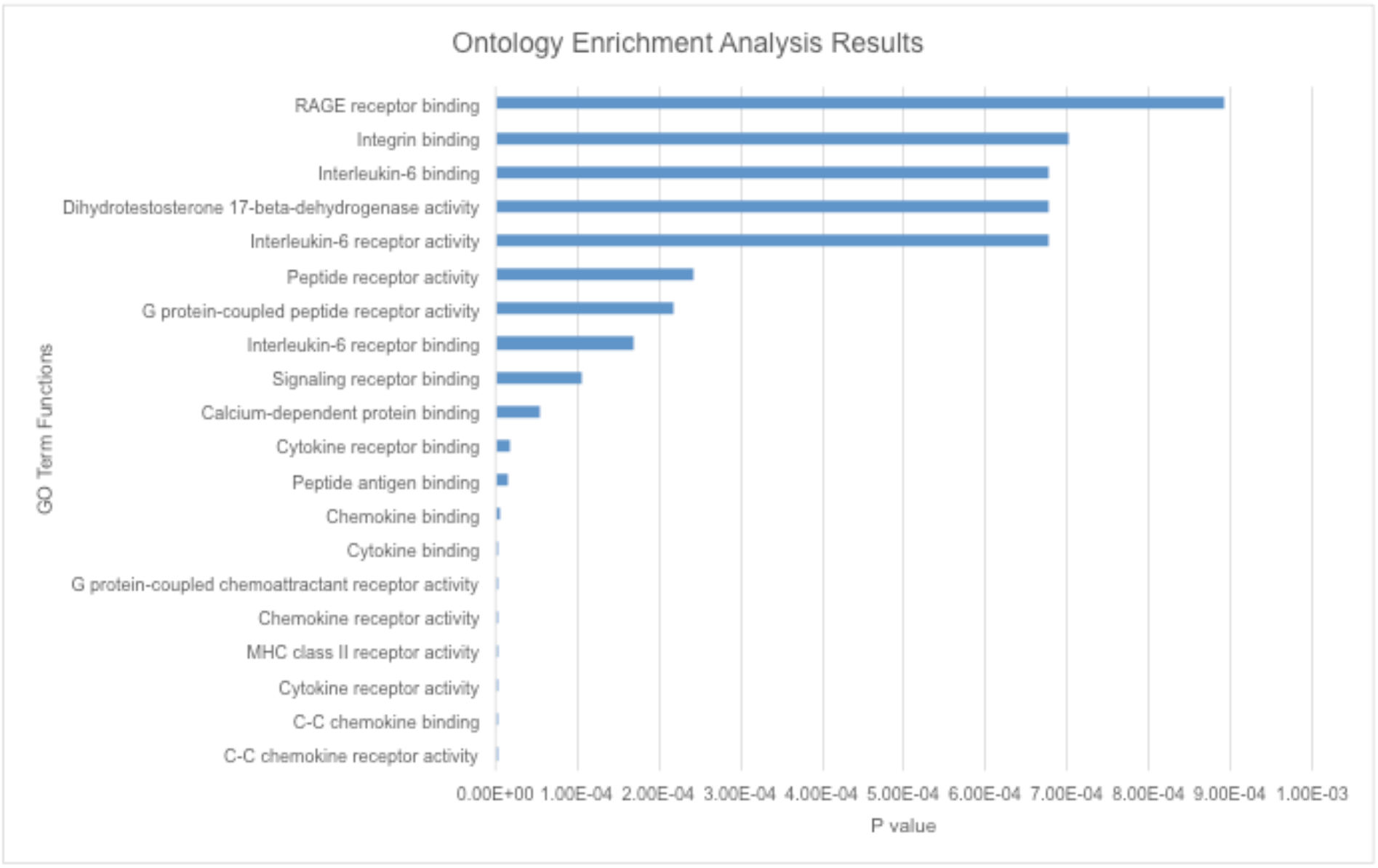
Bar graph summarizing results from gene ontology analysis of genes expressed in human monocytes and encoded within the TADs that encompass the JIA risk LD blocks. These genes are predictably related to basic innate immune functions.

### SNPs within monocyte GROseq peaks are associated with expression levels of genes that are plausible targets of these enhancers

Using BedTools, we identified 12 SNPs, situated in 4 different loci, that were previously identified by MPRA and that also overlapped with monocyte GROseq peaks. Because enhancers typically regulate genes within the same TAD,(25) we posited that the target genes influenced by these SNPs would be one (or more) or those genes within the TADs previously defined. We therefore queried the Genotype-Expression (GTEx) data set using the eQTL calculator and querying whole blood gene expression data, as described in the Methods section. Each of these 12 SNPs was shown to be an eQTL for at least one gene within the monocyte TADs, as shown in **Table 7**.

**Table 7:** eQTL SNPs Within Monocyte GROseq Peaks

| SNP | Locus | Gene | P-Value | P-Value Threshold |
| --- | --- | --- | --- | --- |
| rs4688012 | *TIMMCD1* | *TIMMCD1* | 0.00014 | 0.00023 |
| rs11581043 | *IL6R/ATP8B2* | *ATP8B2* | 2.2e-11 | 0.00018 |
| rs10739578 | *TRAF1* | *C5* | 4.0e-9 | 0.00020 |
|  |  | *PSMD5* | 0.00013 | 0.00021 |
|  |  | *PSMD5-AS1* | 3.4e-36 | 0.00020 |
|  |  | *TRAF1* | 1.8e-11 | 0.00022 |
| rs2109896 | *TRAF1* | *C5* | 1.5e-8 | 0.00020 |
|  |  | *PSMD5* | 0.000018 | 0.00021 |
|  |  | *PSMD5-AS1* | 5.2e-42 | 0.00020 |
| rs7859805 | *TRAF1* | *C5* | 4.1e-8 | 0.00022 |
|  |  | *PSMD5* | 0.000017 | 0.00021 |
|  |  | *PSMD5-AS1* | 4.8e-40 | 0.00020 |
| rs10985080 | *TRAF1* | *C5* | 8.5e-9 | 0.0002 |
|  |  | *PHF19* | 0.000068 | 0.00021 |
|  |  | *PSMD5* | 0.000051 | 0.00021 |
|  |  | *PSMD5-AS1* | 4.5e-43 | 0.00020 |
|  |  | *TRAF1* | 1.3e-8 | 0.00022 |
| rs2549004 | *IRF1* | *C5orf56* | 1.3e-14 | 0.00024 |
|  |  | *SLC22A5* | 9.9e-7 | 0.00028 |
| rs2549007 | *IRF1* | *C5orf56* | 9.3e-15 | 0.00024 |
|  |  | *SLC22A5* | 0.0000012 | 0.00028 |
| rs2549009 | *IRF1* | *C5orf56* | 6.0e-14 | 0.00024 |
|  |  | *SLC22A5* | 1.2e-7 | 0.00028 |
| rs2706385 | *IRF1* | *C5orf56* | 7.5e-16 | 0.00025 |
|  |  | *SLC22A5* | 0.0000023 | 0.00028 |
| rs2706386 | *IRF1* | *C5orf56* | 5.3e-16 | 0.00024 |
|  |  | *SLC22A5* | 0.0000011 | 0.00028 |
| rs41525648 | *IRF1* | *C5orf56* | 0.0000014 | 0.00024 |
|  |  | *SLC22A5* | 4.0e-14 | 0.00028 |

SNP – single nucleotide polymorphism, rs – reference SNP cluster ID

Thus, human data corroborate the in-silico analysis, demonstrating that SNPs within monocyte enhancer regions are associated with alterations in baseline gene expression in whole blood.

## Discussion

We have previously hypothesized that genetic variants of JIA may confer disease risk by modulating both innate and adaptive immune systems(14). We have also shown that characterizing the chromatin architecture of risk haplotypes yields a strategy for elucidating genetic mechanisms, including the identification of genes that are likely to be affected by genetic variants within non-coding regions (the so-called “target genes”). Furthermore, this strategy may assist in identifying the cell types whose functions are impacted by disease-driving variants on risk haplotypes. For example, we have shown that, in immune cells such as neutrophils and CD4+ T cells, the JIA risk haplotypes are enriched for epigenetic features of non-coding regulatory functions, including H3K4me1 and H3K27ac, which commonly identify poised or active enhancers (9-11). Similarly, we have shown that the genetic risk loci for intracranial aneurysm are highly enriched for these same epigenetic features in endothelial cells but not immune cells, which can be observed in affected vessels.(34) Our interest in monocytes/macrophages in the current study extends our interest in the interplay between innate and adaptive immunity (14), which may be a feature of other autoimmune disease such systemic sclerosis/scleroderma (22).

In this study, we investigated the chromatin architecture surrounding 36 JIA risk loci in CD14+ monocytes. We found that these regions, like those in neutrophils and CD4+ T cells, are enriched for epigenetic signatures of non-coding regulatory functions. This was true whether we examined post-translational modifications to histones (H3K4me1/H3K27ac-identified on ChIPseq), or a more functional read-out, the presence of bi-directional RNA synthesis as identified on GROseq (23). We also provide additional evidence that JIA-associated genetic variants may alter enhancer function in these regions: variants that show intrinsic ability to alter gene expression on an MPRA in myeloid K562 cells are located within regions that have a GROseq peak in human macrophages. Using GTEx whole blood expression data, we showed that these SNPs strongly influence the expression of genes within the TADs harboring the GROseq defined enhancers. The GTEx analysis is unlikely to reflect effects exerted in CD14+ cells alone, since they make up only a small part of the signal in whole blood expression profiles. It is just as likely that the observed effect comes from neutrophils, the most abundant cell in the peripheral blood, and, like CD14+ monocytes, of myeloid lineage. However, these findings demonstrate that SNPs within the defined regions are functionally significant.

The enhancer regions we identified in CD14+ monocytes and macrophages are situated within chromatin loops that encompass 321 coding genes that are expressed in these cells. This is a slightly larger number of genes than are situated in the TADs that encompass the JIA haplotypes in CD4+ T cells, where we have identified 287 expressed genes. Thus, JIA-driving risk variants potentially influence a greater number of genes in CD14+ monocytes/macrophages than in CD4+ T cells. Furthermore, the range of cellular functions likely to be impacted by disease-driving variants is broad. Gene ontology analyses of these candidate target genes show significant enrichment for 14 biological different processes, the majority of which are immune related (**Figure 3**).

It is important to note that many of these potential target genes are not situated on the JIA risk haplotypes themselves. The field has become increasingly aware that, for complex traits, genetic effects are not always exerted on the most proximal gene (in terms of linear genomic distance), to the SNPs that are used to tag the genetic risk loci on GWAS or genetic fine mapping studies.(35) Furthermore, Pelikan et al have published direct evidence that enhancer-associated variants on risk haplotypes for systemic lupus erythematosus exert their strongest effects on genes not actually on the risk haplotypes (36). These findings highlight the importance of considering the 3D structure of the genome and chromatin in probing genetic mechanisms that influence JIA risk and/or disease course (8).

While monocytes/macrophages are well-known to be involved in the pathobiology of systemic onset JIA (sJIA) (15, 37-41), less is known about their role in the pathogenesis or pathobiology of the polyarticular/oligoarticular forms of JIA. The most recent understanding stems from alterations to polarization patterns of macrophage activation under JIA inflammatory conditions (15). Circulating monocytes that are recruited to the synovium are stimulated by inflammatory mediators such as TNFα, GM-CSF, IL-1β, and IL-10 to differentiate into tissue resident macrophages. The tissue macrophages are stimulated by the hypoxic environment, microRNAs, and other mediators to differentiate into mixed inflammatory polarizations. Cytokines, chemokines, and growth factors such as VEGF that have been shown to be upregulated in the synovial fluid and serum of patients with JIA are secreted by these synovial monocytes and macrophages. Furthermore, the products of these monocyte/macrophages are the targets of several therapeutics used to treat JIA, such as TNFα inhibitors etanercept and adalimumab, IL-1β receptor antibody anakinra, and IL-6 receptor antibody tocilizumab (15). Our findings support the idea that many of these steps may be under genetic influence and that genetically-mediated variation in these processes may contribute to disease risk.

Although inflammatory states may recruit circulating monocytes to enter tissue and differentiate into tissue resident macrophages, the circulating monocytes only contribute to forming small portion of the permanent population of tissue macrophages that remain after the inflammatory period has concluded (42). Tissue resident macrophages may instead have origins from variable populations, including embryonic yolk-sac precursors and fetal liver derived hemopoietic stem cells, in addition to adult monocytes (42). This lack of consistency and shared lineage in the cells that repopulate tissue resident macrophages suggests that genetic risk variants might occur in the more cohesive population of tissue resident macrophages themselves rather than their diverse monocyte precursors or circulating CD14+ monocytes/macrophages.

There are several other limitations to this study that need to be considered. The first is that the presence of H3K4me1 and/or H3K27ac ChIPseq peaks, even in a region of open chromatin, is not *prima facie* evidence that a region has enhancer activity. Invariably, enhancer activity has to be confirmed experimentally, preferably in the context of native chromatin. In contrast, the presence of bidirectional RNA synthesis, as detected on GROseq, provides a strong functional read-out of enhancer activities, as the synthesized strands are invariably so-called enhancers RNA (eRNA) that facilitate the interaction between the enhancer and promoter (23).

It is also important to note that not every patient with JIA has a disease-driving variant on every haplotype. Thus, although we can make inferences as to the spectrum of innate immune functions that might be altered in patients harboring variants on the risk haplotypes (e.g., **Figure 3**), this does not mean that these pathways are affected in every patient. Indeed, the impetus toward precision medicine initiatives is based on the premise that each patient’s genetic vulnerabilities (and strengths) is unique, and that therapy should be directed to those unique features of an individual patient’s disease biology.

In conclusion, we provide evidence, based on the chromatin architecture surrounding the genetic risk haplotypes, that genetic risk for JIA exerts effects in cells of the monocyte-macrophage lineage. These findings provide a strong rationale to test this concept empirically, using both patient cells(36) and in vitro approaches (43).

## Acknowledgments

The authors wish to thank Drs. John Lis and Charles Danko for their guidance in analyzing and interpreting GROseq data using their dReg pipeline.

## Author contributions

EAC – Performed data analyses and interpretation. Assisted in writing the manuscript.

EKH – Performed data analysis and interpretation. Assisted in writing the manuscript.

KEP – Developed analysis pipelines and methods, assisted in data analysis.

KJ – Assisted in data analysis, interpretation, and preparation of the manuscript

VMT - Developed analysis pipelines and methods, assisted in data analysis.

JNJ – Designed and directed the study, assisted with data analysis and interpretation, and assisted with preparation of the manuscript.

---

**Supplemental Table:**
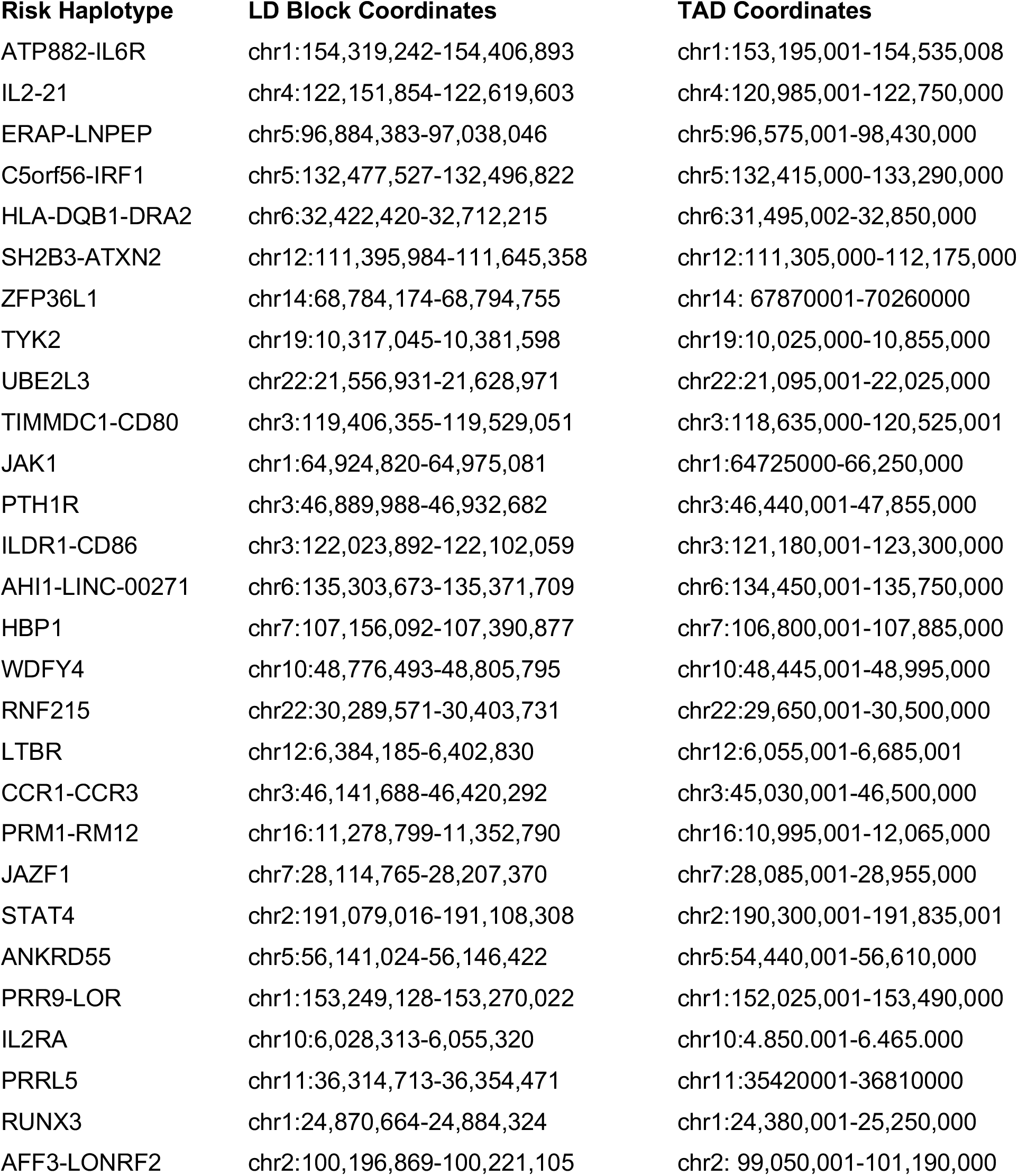
Genomic locations of TADs surrounding 28 JIA risk haplotypes

